# Testing the redox theory of aging under parasitism

**DOI:** 10.64898/2025.12.03.692220

**Authors:** Luís M. Silva, Alessandro Belli, Jacob C. Koella

## Abstract

The redox theory of aging proposes that an oxidative imbalance, possibly amplified by infection, drives senescence. We experimentally evolved mosquitoes under early or late reproduction with or without parasite exposure, and quantified longevity, fecundity, and redox markers. Although selection generated the expected life-history divergence, there was only a poor non-linear association between a redox gradient and longevity. Thus, oxidative stress contributed to, but did not determine, the evolution of aging.

## Introduction

Aging evolves because the force of natural selection declines with age ^1–4^, allowing late-acting deleterious effects to accumulate ^5^. As a consequence, evolutionary theories, such as the *selective shadow* (or antagonistic pleiotropy) and the *disposable soma hypothesis* ^2,5,6^, predict a trade-off between investing in early reproduction and maintaining somatic function later in life. Under this framework, Rose and colleagues (and subsequent research groups) demonstrated that selection for early reproduction should favour high fecundity at the expense of longevity ^7–9^.

The physiological mechanisms, however, through which these life history trade-offs shape aging remain under active debate ^10,11^. One well-known hypothesis is the *redox theory of aging* (RTA) ^12^. The latter proposes that age-related decline results from shifts in the balance between reactive oxygen species (ROS), antioxidant defences, and cellular repair. Therefore, life-history traits associated with high reproductive effort or chronic immune activation should push organisms toward more oxidized physiological states, and life histories that favour late reproduction should promote stronger antioxidant capacity. However, empirical tests that explicitly link long-term selection on life history with changes in oxidative physiology and aging are rare.

Complicating these ideas, organisms in nature face frequent and even chronic immune challenges that affect reproductive investment, metabolic allocation, and redox processes ^13–15^ and thus shape how aging evolves. Infection is often expected to accelerate physiological decline, either through direct pathology (immunopathology) ^16^, oxidative immune responses (*e.g.,* ROS and inflammaging) ^17^, or energetic diversion away from somatic maintenance. Yet, hosts can also adopt alternative strategies, such as disease tolerance ^18^, that mitigate fitness costs without reducing the parasite burden. Whether parasitism restructures redox physiology or interacts with life-history evolution to alter aging remains largely unresolved.

Hence, we set out to test the RTA in a mosquito-microsporidian system. In particular, we examined whether the pressure of parasitism disrupts the redox balance and its evolution. We experimentally selected *Aedes aegypti* females to reproduce either early (7–14 days after emergence) or late (35–42 days) either with or without exposure to their natural microsporidian parasite, *Vavraia culicis* ^19^, which triggers oxidative immune responses ^13^. After at least ten generations, we quantified how selection on reproduction and parasitism shaped life history **(Fig. 1)** and redox expression **(Fig. 2)**. More broadly, we asked whether the RTA can be applied to ecological and epidemiological settings, where hosts are continually exposed to parasites and must navigate the inherent trade-offs that shape the evolution of aging.

**Figure 1.**
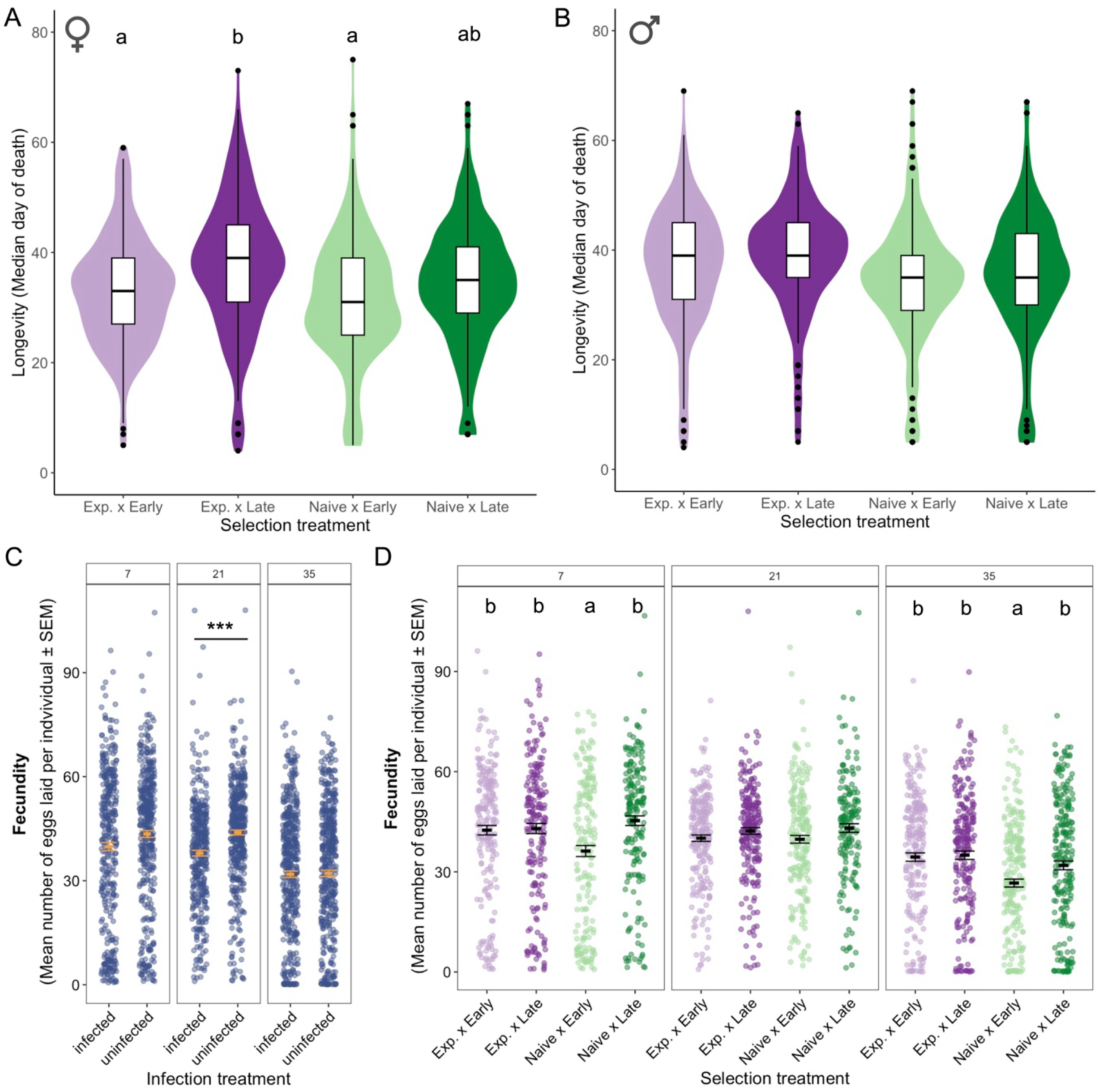
Evolution of longevity and fecundity across selection treatments. Median longevity of female **(A)** and male **(B)** mosquitoes after 10 generations of selection for early or late reproduction, with or without exposure to the parasite *V. culicis*. Sample sizes were: 315 females and 284 males in Exposed and Early; 308 females and 287 males in Exposed and Late; 281 females and 274 males in Naive and Early; and 295 females and 300 males in Naive and Late. Effect of **(C)** infection and **(D)** selection on fecundity, measured at three ages (7, 21 and 35 days after emergence). Each point is an individual, and the black bars show means and the respective standard errors. Asterisks (***) indicate a significant difference, with a *p*-value of < 0.001, whereas letters denote results of post-hoc multiple comparisons tests conducted after the respective GLMM. Absence of letters indicates no difference between treatments, according to the multiple comparisons test. Sample sizes for the fecundity assay in the presence or absence of infection were 359 and 404 females at day 7, 371 and 442 at day 21, and 470 and 471 at day 35, respectively. Across selection treatments, sample sizes at day 7 were 203 (Exposed–Early), 194 (Exposed–Late), 184 (Naive–Early), and 182 (Naive–Late); at day 21 they were 223, 222, 205, and 153; and at day 35 they were 237, 232, 233, and 239, respectively.

**Figure 2.**
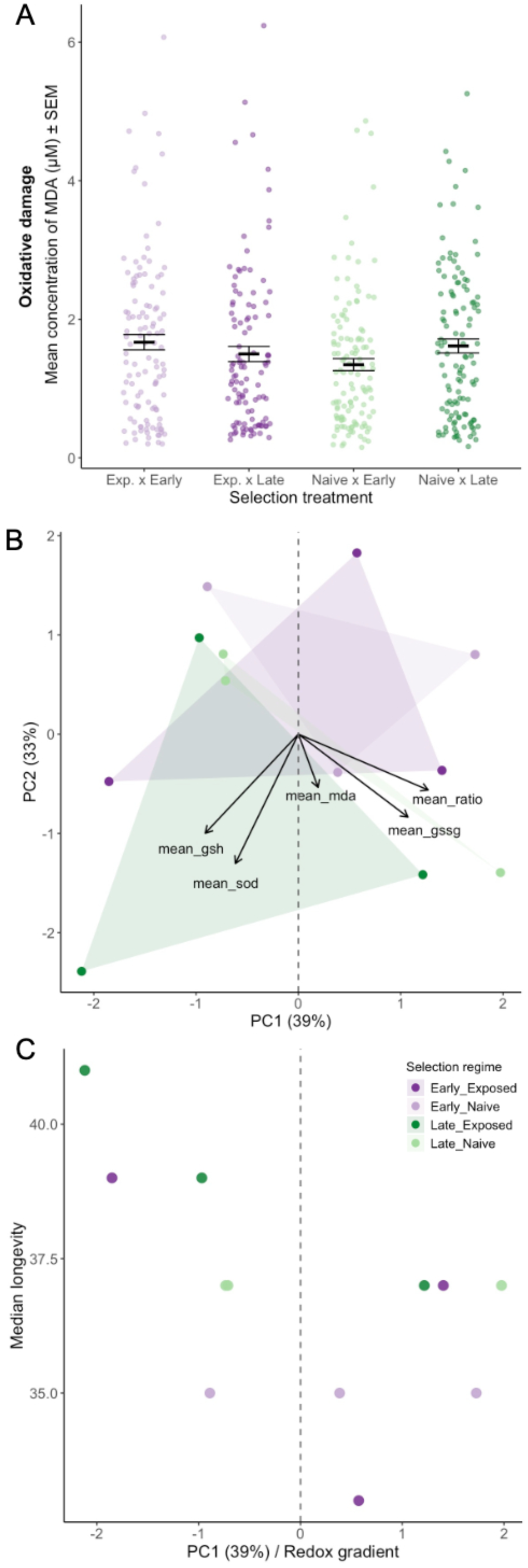
Evolution of oxidative balance and aging. Oxidative balance of selected individuals was measured by assessing the concentrations of malondialdehyde (MDA - marker of oxidative damage), superoxide dismutase (SOD - marker of antioxidant activity), and individual values, as well as the ratio between oxidized and reduced glutathione (GSH: GSSG - marker of oxidative balance) without infection. **(A)** Only MDA was found to be affected by parasite- and reproduction-based selection (independently), although marginally and therefore not detectable in post hoc multiple comparisons tests. Sample size consisted of 117 “Exposed-Early”, 109 “Late-Exposed”, 122 “Naive-Early” and 120 “Naive-Late” females. **(B)** PCA of oxidative markers. PC1 (39% variance) captures a gradient from oxidized to reduced redox state. Negative loadings are associated with antioxidant capacity (GSH and SOD), whereas positive loadings are associated with oxidative damage (MDA, GSSG, and GSSG: GSH ratio). Arrows indicate marker loadings. **(C)** Relationship between redox PC1 and median adult longevity across replicates of the different selection treatments.

## Results

Experimental evolution **(Supplementary Fig. 1)** generated a divergence in aging and reproduction among the four selection regimes **(Fig. 1)**. Here, *exposed* and *naïve* denote the evolutionary selection history, whereas *infected* and *uninfected* denote the post-selection assay treatment. In this host–parasite association, *V. culicis* causes relatively low virulence in *A. aegypti*, such that evolutionary exposure represents a chronic, low-pathology regime. Females selected for late reproduction lived significantly longer than those selected for early reproduction (*χ²* = 20.62, *df* = 1, *p* < 0.001; **Fig. 1a**), but males did not **(Fig. 1b),** as expected from evolutionary predictions. Parasite exposure increased host longevity in both sexes (Females: *χ²* = 1.29, *df* = 1, *p* = 0.005; Males: *χ²* = 1.29, *df* = 1, *p* = 0.040; **Fig. 1; Supplementary Table 1**). However, upon the death of the mosquitoes, selection treatments did not differ in the proportion of clearance or in the parasite load **(Supplementary Fig. 2)**. In other words, lines with a history of parasite exposure evolved greater survival despite no evidence for increased resistance (no differences among selection regimes in parasite clearance or parasite load at death), consistent with an evolutionary shift toward disease tolerance rather than enhanced parasite control.

We quantified five markers of redox balance: malondialdehyde (MDA) as a measure of oxidative damage; superoxide dismutase (SOD) as an indicator of antioxidant activity; and the glutathione ratio and its components (GSH and GSSG) as markers of oxidative balance. Among these, only MDA showed a marginal interaction between reproductive regime and parasite exposure (*χ²* = 4.63, *df* = 1, *p* = 0.031; **Fig. 2a; Supplementary Fig. 3**). Oxidative damage differed subtly across treatments, with parasite exposure slightly increasing MDA levels in early-reproducing lines but not in late-reproducing lines. This interaction was weak and highly overlapping among groups, indicating that neither factor produced strong or consistent divergence on its own. Overall, although parasitism modestly modulated MDA depending on reproductive regime (*χ²* = 5.17, *df* = 1, *p* = 0.023), the evolved lines exhibited broadly similar oxidative damage profiles across treatments **(Fig. 2a)**.

Nevertheless, we observed significant variation in the redox markers. Hence, to summarize covariation among markers, we performed a principal component analysis (PCA) on replicate-level means. PC1 explained 39% of the variance and captured an oxidative–reductive gradient: higher scores reflected more oxidized profiles with elevated MDA, GSSG and GSH:GSSG ratio, whereas lower scores reflected more reduced profiles with higher GSH and SOD **(Fig. 2b, Supplementary Tables 3 and 4**). PC2 explained a further 33% of the variance and represented residual differences among markers. Lines selected for early reproduction, especially under parasite exposure, tended to occupy the more oxidized end of PC1, while late-selected lines were generally more reduced. However, all four selection regimes still overlapped in redox space, indicating that evolutionary responses in oxidative physiology were modest and heterogeneous.

We then asked whether these multivariate redox profiles were associated with evolved differences in aging. Across lines, PC1 scores explained a moderate fraction of the variation in median longevity of uninfected females **(Fig. 2c, Supplementary Table 5)**. A linear mixed model (LMM) indicated that more reduced profiles tended to be longer-lived. A generalized additive model (GAM) showed a weak but significant non-linear association between PC1 and median longevity (s(PC1): edf = 2.2, *F* = 5.0, *p* = 0.026), consistent with redox state explaining only a fraction of the among-line variation in lifespan. Thus, the baseline oxidative state was related to longevity, but not in the simple, canonical form often envisioned by the redox theory of aging. Overall, both univariate and multivariate analyses support a role for redox physiology in shaping aging; however, they also indicate that it only partially accounts for the substantial divergence in life histories generated by experimental evolution.

## Discussion

Experimental evolution in our system revealed the life-history trade-offs predicted by theories of the evolution of senescence, but their mechanistic basis cannot be explained by the RTA alone. Selection for early reproduction resulted in short-lived mosquitoes and a shift toward earlier reproductive output, whereas selection for late reproduction extended lifespan, consistent with evolutionary predictions of the selective shadow (or antagonistic pleiotropy) ^5^ and the disposable soma hypothesis ^6^. Consistent with this, fecundity differences were age-dependent rather than uniformly elevated in early-selected lines. Together, these findings highlight two key points: (i) aging cannot be solely explained by shifts in oxidative balance; and (ii) biology operates as an evolutionary game played on a life-history board ^20^, where aging emerges from strategic allocation between reproduction and somatic maintenance and traits ^21^. Our data also aligns with principles described by Charlesworth (1994), which state that the tempo of aging is governed by the age-specific strength of selection, rather than simply by mechanistic deterioration^4^. Hence, aging responds to how selection is distributed across the life cycle, and to life-history trade-offs between reproduction, survival, and defence, rather than to damage accumulation alone. Because traits were measured in different individuals, our results address evolved trade-offs among selection regimes, not within-individual allocation decisions.

The redox theory of aging provides a robust mechanistic framework by linking oxidative balance to age-related decline. In our experiment, however, redox physiology responded more weakly and less coherently than life history. Baseline oxidative markers differed only slightly among treatments, and the main multivariate redox axis (PC1) did not separate the four selection regimes. Nevertheless, lines at the more reduced end of the PC1-axis tended to live longer, whereas more oxidized or intermediate profiles were generally shorter-lived. The non-linear form of this relationship and the broad overlap among treatments in redox space suggest that a simple one-to-one link between oxidative state and the pace of aging is unlikely. Instead, they suggest that redox is one of several mechanistic pathways by which life-history evolution can influence senescence. This partial decoupling is consistent with critiques that oxidative biomarkers often fail to predict lifespan evolution across multiple taxa^22^. After all, ROS has been broadly recognized as functioning not only as damaging oxidants but also as integral signalling molecules regulating immunity, metabolism, and mitochondrial dynamics^23^.

The effects of chronic exposure to parasitism further illustrate this complexity. Contrary to the expectation that infection accelerates physiological decline ^24^, parasite exposure increased longevity in both sexes. Importantly, the parasite used here exhibits relatively low virulence in *A. aegypti* under our conditions ^14,25^. Thus, our conclusions primarily address how chronic low-virulence exposure interacts with life-history evolution and redox physiology. Under stronger parasite pressure (*e.g.,* higher virulence, higher doses, or parasites causing greater tissue damage), selection could instead favour increased resistance, stronger oxidative immune responses, and potentially a tighter coupling between redox state and survival. Evolving with parasites did not increase the ability to clear parasites or reduce parasite load. This pattern is consistent with evolutionary shifts in resource allocation and damage management, including changes in disease tolerance (the ability to mitigate infection-associated costs on host fitness) ^18,26,27^, rather than a straightforward enhancement of oxidative defences. Tolerance is expected to evolve when clearance is costly, ineffective, or the parasite does not pose a sufficient danger that necessitates its elimination ^18^. Our parasite, *V. culicis*, causes relatively low virulence in mosquitoes, particularly in *A. aegypti* ^25^, although it still triggers an immune response from its hosts ^28^. Hence, a low-virulence parasite, combined with constraints from early *versus* late reproduction, may favour disease tolerance rather than stronger resistance, particularly when resistance to this parasite already appears significant ^25^. Allowing mosquitoes to evolve for many generations under these joint pressures, rather than relying on short-term manipulations or single-generation correlations, we capture this evolutionary realism. It reveals that redox leaves only a modest, context-dependent imprint, even when aging itself evolves strongly.

These findings have broader implications for the biology of aging. First, they emphasize that aging should be studied in a multidimensional space, which encompasses several traits regulated by interactions between metabolic allocation, immune activation, endocrine pathways, mitochondrial function, and even behavioural ecology. This multidimensionality reflects the “hallmarks of aging”, which emphasize that molecular and physiological pathways (from nutrient sensing to mitochondrial dynamics) interact to shape lifespan ^29^. Redox processes contribute to these dynamics, but they do not operate in isolation. Second, the evolution of tolerance in our system highlights how the expression of this specific strategy might be especially advantageous in the wild. An increase in tolerance in a host population is likely to be followed by a rise in the prevalence of the parasite ^18^, which is amplified by increases in disease transmission among infected tolerant hosts ^30^. This has severe consequences for aging, as well as host-parasite coevolution and epidemiology.

Ultimately, our findings underscore the importance of integrating mechanistic aging research with an ecological and evolutionary perspective. Organisms rarely live in benign laboratory environments; instead, they face sustained immune challenges and fluctuating reproductive opportunities that shape the evolution of aging. Through the combination of experimental evolution with multivariate physiological analysis, we demonstrate that lifespan can evolve robustly even when canonical oxidative markers shift only slightly. A deeper understanding of how aging mechanisms intersect will be essential for predicting the evolution of aging and for designing evolutionarily informed approaches to promote healthy aging and improve age-specific responses to infection.

## Materials and Methods

### Experimental model

We used the microsporidian parasite *Vavraia culicis floridensis,* provided by James Becnel (USDA, Gainesville, FL) and the UGAL strain of the mosquito *Aedes aegypti* (obtained from Patrick Guérin, University of Neuchâtel). Mosquitoes were reared under standard laboratory conditions (26 ± 1°C, 70 ± 5% relative humidity, and 12 h light/dark). *V. culicis* is a generalist, obligatory, intracellular parasite that infects mosquito epithelial gut cells. Mosquitoes are exposed to the microsporidian during the larval aquatic stage, where they ingest spores along with their food in the water. After replication within the larvae, as adults, they start producing spores from within the gut cells, re-infecting new cells and propagating the cycle until they kill the host. Spores are transmitted when infected individuals die in water or from mother to offspring by adhering to the surface of the eggs. Before starting the experiment, *V. culicis* was kept in large numbers, alternating every generation between infecting *A. aegypti* and *Anopheles gambiae* populations to ensure it remains a generalist parasite. The infection protocol was performed as described in detail in the experiments below, with two-day-old larvae being exposed to 10,000 spores per larva. In brief, deceased individuals were collected every other day, placed into 2 mL Eppendorf tubes containing 1 mL of deionized water and a stainless-steel bead (Ø 5 mm), and stored at 4℃. These individuals (approximately 15 per tube) were later homogenized using a Qiagen TissueLyser LT at a frequency of 30 Hz for 2 minutes and then used to infect new larvae.

### Experimental evolution and rearing

We performed experimental evolution to create four selection treatments in a fully factorial design: selection for early *vs*. late reproduction, and selection in the absence *vs*. presence of *V. culicis*. Each regime was replicated four times (16 lines total). In every generation of selection and maintenance, we ensured a density of at least 200 adult individuals. To start the experiment, eggs from the stock UGAL colony were hatched synchronously under reduced air pressure and distributed among 16 lines. Each line was maintained separately across ten generations with a standardized density of larvae per generation. Larvae were reared in deionized water (800 ml in a plastic tray) and fed daily with Tetramin Baby ® fish food in the following daily doses: 0.04, 0.053, 0.107, 0.213, 0.64, 0.32 mg/larva for days 1, 2, 3, 4, 5, 6 and later, respectively. At the pupae stage, individuals were transferred to cups (5 cm Ø x 10 cm) placed inside cages (30 x 30 x 30 cm), and adults were provided with *ad libitum* access to 6% sucrose.

For selection lines exposed to the parasite, larvae were exposed to *V. culicis* spores 48 h after hatching by adding 20,000 spores/larva (infective stage) to the water and gently mixing. Naive (control) lines experienced identical handling but received parasite-free water. Spore concentration was assessed using a hemocytometer under a phase-contrast microscope before each infection.

All females received their first blood meal at a fixed age interval post-hatching. Females were fed on human skin (AB’s and LMS’ arm) for selection and post-selection readouts. In early-reproduction lines, only eggs laid in the first clutch were used to found the next generation. In late-reproduction lines, females received repeated blood meals at fixed intervals, and only eggs from the last clutch (collected when approximately two-thirds of females had died) were used to start the following generation. This design generated four selection treatments: (i) exposed x early; (ii) exposed x late; (iii) naive x early; and (iv) naive x late. Before measuring responses to selection, a generation without any selective pressures was given to avoid parental effects on our post-selection measurements.

### Responses to selection

Before assessing any trait, selection was halted for one generation to avoid carry-over effects from parental generations. Longevity and oxidative stress were assessed at generation F12, while fecundity was assessed at F18. Assays were performed at different generations for logistical reasons and because longevity assays are long-running and space-intensive. Longevity and redox physiology were quantified at F12, whereas fecundity was quantified later (F18). Importantly, lines were continuously maintained under the same selection regimes between these readouts, such that the selected life-history differences were preserved. Moreover, before each assay, we included a common-garden generation without selection to reduce parental or carry-over effects. Because our primary aim is to quantify evolved differences among selection regimes and replicate lines, rather than temporal dynamics across generations, measuring traits at different post-selection generations does not change the qualitative evolutionary contrasts tested here.

Longevity, fecundity, and redox markers were measured in different individuals. This was necessary for two reasons. First, longevity assays are terminal and preclude repeated measurements of reproduction later in life. Second, quantifying redox markers requires destructive sampling, which cannot be combined with subsequent measurements of fecundity or survival in the same mosquito. Therefore, our inference about trade-offs is evolutionary and population-level: we test whether selection regimes that differ in lifespan and reproductive allocation also differ in redox physiology across replicate lines, rather than measuring within-individual allocation decisions. Across assays, cohorts were generated from the same selection lines and reared under common standardized laboratory conditions. Infection treatments (where applied) were administered at the same larval age and dose. Thus, while endpoints were measured in different individuals, comparisons are made within a controlled common environment, and treatment effects are evaluated at the line/regime level, which is the unit that evolved during selection.

#### Longevity assay

To quantify evolved differences in lifespan and infection outcomes, we reared larvae from each selection line individually. Eggs from each line were synchronously hatched, and 48-hour-old larvae were transferred and individualized in single wells of 12-well plates, containing 3 mL of deionized water, and fed daily as above. For each line, half of the larvae were infected with *V. culicis* (20,000 spores per larva) and half remained uninfected. Pupae were then transferred individually to small cups enclosed within larger cages, and adults had ad libitum access to 6% sucrose. Mosquitoes were monitored daily until death. Dead individuals were frozen for later assessment of infection status and parasite load by homogenizing whole bodies in deionized water and counting spores with a hemocytometer. These data were used to quantify longevity, infection probability, and parasite burden across selection regimes.

#### Fecundity assay

To quantify the effect of selection on reproduction, we measured fecundity in all selection lines in a full-factorial design (four selection treatments x four replicate lines x two infection treatments [infected or not]). Freshly emerged larvae (3–6 h old) from each line were reared individually in 24-well plates (one larva per well, 3 mL deionized water per well) and fed daily as described above. Larvae assigned to the infected group were exposed to 20,000 spores in 3 mL of water at 2 days post-hatching. In contrast, uninfected controls received parasite-free water. Pupae were transferred to 180 mL cups within 30 × 30 × 30 cm cages. To control for paternal effects, males emerging from the selection lines were replaced by stock males at a 1:1 male-to-female ratio. Adults were allowed to mate for seven days in groups of 100 per cage, after which stock males were removed. Females were offered a human blood meal (LMS arm) for 7–8 minutes at three adult ages: 7-, 21-, and 35 days post-emergence. Two days after each blood meal, females were gently transferred to individual oviposition cups (standard 180 mL plastic cups with moist filter paper) and kept there for egg laying. Females were then frozen at −20 °C. We counted the number of eggs per female per time window, and measured wing length as a proxy for body size.

#### Oxidative stress assay

To measure evolved differences in redox physiology, we quantified oxidative stress markers in uninfected females from three (out of four) randomly selected replicates of each selection treatment. For each line, 120 larvae were hatched under reduced air pressure and reared individually in 12-well plates (3 mL deionized water per well). Pupae were transferred to 180 mL cups for emergence, and adults were kept in 30 x 30 x 30 cm cages with 6% sucrose solution. Females were collected at specific ages (5- and 15-days post-emergence), killed by freezing at −80 °C, and stored individually in 2 mL microcentrifuge tubes. For each mosquito, the wings and legs were removed, and the body was weighed to the nearest 0.01 mg. The body was then homogenized in 120 µL phosphate-buffered saline using a 2 mL tube with a 5 mm stainless-steel bead in a Tissue Lyser (4 min, 45 Hz). Homogenates were centrifuged (10,000 rpm, 10 min, 4 °C), and supernatants were split into aliquots for glutathione (GSH, GSSG), MDA, and SOD assays. GSH and GSSG were extracted and quantified by UHPLC–MS/MS using established protocols ^31^. MDA was measured by HPLC–MS against a standard curve prepared from MDA tetrabutylammonium salt, after a phase-partitioning and evaporation step. SOD activity was measured using the Superoxide Dismutase Assay Kit (Cayman Chemical), with mosquito homogenates diluted 1:8 in PBS; samples were run in duplicate and averaged for analysis. There was no difference across selection treatments in either time point, so they were combined in the analyses and figures of the study.

### Statistical analyses

All analyses and plots were performed with R version 4.5.2 in RStudio version 2025.09.2+418 using the packages “car”, “lme4”, “DHARMa”, “ggplot2”, “dplyr”, “tidyr”, “emmeans”, “multcomp”, “scales”, “purrr”, “mgcv” and “mgcViz”. https://Biorender.com was used to assemble the figure panels. Significance was assessed with the “Anova” function of the “car” package. We used a Type III ANOVA when a significant interaction was present, and a Type II ANOVA otherwise. When relevant, we performed post hoc multiple comparisons using the “emmeans” package with the default Tukey adjustment.

Longevity (median age at death) was analyzed using a linear mixed model (LMM) with reproductive regime (early *vs.* late), parasite exposure during evolution (naïve *vs.* exposed), and their interactions as fixed effects, and line as a random effect. This was performed separately for each sex **(Supplementary Table 1)**.

Fecundity (egg counts of egg-laying females transformed by a quadratic function) was analyzed using an LMM with reproductive selection, parasite selection, and infection as explanatory variables, as well as their interaction, and replicate as a random factor. A separate LMM was run for each fecundity window **(Supplementary Table 2)**.

Redox markers (GSH, GSSG, GSH: GSSG ratio, MDA, SOD) were also analyzed using LMMs with reproductive regime, parasite exposure during evolution, and their interactions as fixed effects, and line as a random effect. To assess changes in redox physiology across selection treatments, we performed a principal component analysis (PCA) on the means of each line for the five oxidative markers. Before running the PCA, markers were mean-centred and scaled to unit variance. PCA was conducted using the “prcomp” function in R, and loadings were extracted to identify the main axes of variation **(Supplementary Tables 3 and 4)**. We focused on PC1 and PC2 for visualization and assessed separation among selection regimes qualitatively, given the limited number of replicate lines.

To test whether multivariate redox profiles predicted evolved differences in aging, we related PC1 (redox gradient) scores to the median adult longevity of each line. We first fitted a simple linear model with PC1 as a predictor. We then used a generalized additive model (GAM) with a smooth term for PC1 to allow for non-linear relationships (using the “gam” function in the “mgcv” package) **(Supplementary Table 5)**. Model residuals were inspected to confirm assumptions. Given the small number of lines, we interpreted these analyses cautiously, focusing on the direction and shape of the association rather than on precise effect size estimates.

## Data availability

All data generated are provided as supplementary files

## Code availability

The R script used for statistical analyses is provided as supplementary files.

## Acknowledgments

We would like to thank Alexei A. Maklakov, Alfonso Rojas-Mora, Tiago G. Zeferino, and Thomas Flatt for their valuable feedback throughout the project.

## Author contributions

AB and JCK conceived the project. AB, LMS and JCK designed the experiments. AB and LMS conducted the experiments. LMS analyzed the data and wrote the first draft of the manuscript.

## References

1. Williams, G. C. Natural selection, the costs of reproduction, and a refinement of Lack’s principle. Am. Nat. 100, 687–690 (1966).

2. Hamilton, W. D. The moulding of senescence by natural selection. J. Theor. Biol. 12, 12–45 (1966).

3. Kirkwood, T. B. L. Evolution of ageing. Mech. Ageing Dev. 123, 737–745 (2002).

4. Charlesworth, B. Evolution in age-structured populations. (1994).

5. Rose, M. R. Antagonistic pleiotropy, dominance, and genetic variation. Heredity (Edinb). 48, 63–78 (1982).

6. Lorenzini, A., Stamato, T. & Sell, C. The disposable soma theory revisited: time as a resource in the theories of aging. Cell Cycle 10, 3853–3856 (2011).

7. Rose, M. R. %J E. Laboratory evolution of postponed senescence in Drosophila melanogaster. 38, 1004–1010 (1984).

8. Kirkwood, T. B. L. & Rose, M. R. Evolution of senescence: late survival sacrificed for reproduction. Philos. Trans. R. Soc. London. Ser. B Biol. Sci. 332, 15–24 (1991).

9. Fabian, D. K. et al. Evolution of longevity improves immunity in Drosophila. Evol. Lett. (2018).

10. Slade, L., Etheridge, T. & Szewczyk, N. J. Consolidating multiple evolutionary theories of ageing suggests a need for new approaches to study genetic contributions to ageing decline. Ageing Res. Rev. 100, 102456 (2024).

11. Aronoff, J. E. & Trumble, B. C. An evolutionary medicine and life history perspective on aging and disease: Trade-offs, hyperfunction, and mismatch. Evol. Med. Public Heal. 13, 111–124 (2025).

12. Harman, D. Free radical theory of aging: history. Free radicals aging 1–10 (1992).

13. Silva, L. M. The transcriptome of the mosquito host Anopheles gambiae upon infection by different selected lines of the microsporidian parasite Vavraia culicis. bioRxiv 2009–2024 (2024). 10.1101/2024.09.22.613703

14. Silva, L. M. & Koella, J. C. Complex interactions in the life cycle of a simple parasite shape the evolution of virulence. PLoS Pathog. (2025). 10.1371/journal.ppat.1013294

15. Zeferino, T. G., Rojas Mora, A., Vallat, A. & Koella, J. C. Dietary shifts in infected mosquitoes suggest a form of self-medication despite benefits in uninfected individuals. Proc. B 292, 20251659 (2025).

16. Graham, A. L., Allen, J. E. & Read, A. F. Evolutionary causes and consequences of immunopathology. Annu. Rev. Ecol. Evol. Syst. 36, 373–397 (2005).

17. Baylis, D., Bartlett, D. B., Patel, H. P. & Roberts, H. C. Understanding how we age: insights into inflammaging. Longev. Heal. 2, 8 (2013).

18. Schneider, D. S. & Ayres, J. S. Two ways to survive infection: what resistance and tolerance can teach us about treating infectious diseases. Nat. Rev. Immunol. 8, 889 (2008).

19. Silva, L. M., Zeferino, T. G. & Koella, J. C. Vavraia culicis. Trends Parasitol. (2025). 10.1016/j.pt.2025.02.011

20. Stearns, S. C. The evolution of life histories. 249, (Oxford university press Oxford, 1992).

21. Crews, D. E. & Bogin, B. Growth, development, senescence, and aging: a life history perspective. A companion to Biol. Anthropol. 124–152 (2010).

22. Speakman, J. R. & Selman, C. The free-radical damage theory: accumulating evidence against a simple link of oxidative stress to ageing and lifespan. Bioessays 33, 255–259 (2011).

23. Schieber, M. & Chandel, N. S. ROS function in redox signaling and oxidative stress. Curr. Biol. 24, R453–R462 (2014).

24. Quiros-Roldan, E., Sottini, A., Natali, P. G. & Imberti, L. The impact of immune system aging on infectious diseases. Microorganisms 12, 775 (2024).

25. Zeferino, T. G. & Koella, J. C. Host-specific effects of a generalist parasite of mosquitoes. Sci. Rep. 14, 18365 (2024).

26. Zeller, M. & Koella, J. C. The Role of the Environment in the Evolution of Tolerance and Resistance to a Pathogen. Am. Nat. 190, 389–397 (2017).

27. Akyaw, P. A., Paulo, T. F., Lafuente, E. & Sucena, É. Pathogen-induced damage in Drosophila: uncoupling disease tolerance from resistance. PLoS Pathog. 21, e1013482 (2025).

28. Desjardins, C. A. et al. Contrasting host–pathogen interactions and genome evolution in two generalist and specialist microsporidian pathogens of mosquitoes. Nat. Commun. 6, 7121 (2015).

29. López-Otín, C., Blasco, M. A., Partridge, L., Serrano, M. & Kroemer, G. Hallmarks of aging: An expanding universe. Cell 186, 243–278 (2023).

30. Silva, L. M., King, K. C. & Koella, J. C. Dissecting transmission to understand parasite evolution. PLoS Pathog. 21, e1012964 (2025).

31. Mora, A. R., Firth, A., Blareau, S., Vallat, A. & Helfenstein, F. Oxidative stress affects sperm performance and ejaculate redox status in subordinate house sparrows. J. Exp. Biol. 220, 2577–2588 (2017).

