## Supplementary material for "Testing the redox theory of aging under parasitism": SI

**Supplementary File 1:**
Supplementary tables and figures for "Testing the redox theory of aging under parasitism"
by Luís M. Silva *, Alessandro Belli * and Jacob C. Koella
*These authors contributed equally to this study.

This document includes the following:
Supplementary Table 1. Longevity and fecundity after selection.
Supplementary Table 2. MDA concentration across selection treatments.
Supplementary Table 3. Summary of the principal component analysis.
Supplementary Table 4. Loadings of oxidative markers on PCA.
Supplementary Table 5. Relationship between oxidative PCA axis and longevity.

Supplementary Figure 1. Design of the experimental evolution setup.
Supplementary Figure 2. Post selection measurements: parasite load at death.
Supplementary Figure 3. Post selection measurements: oxidative markers.

#### Tables

**Supplementary Table 1. Longevity and fecundity after selection.** After selecting for early or late reproduction, regardless of exposure to parasitism or not, we measured longevity and fecundity throughout the mosquito's life. First, we assessed changes in longevity in the absence of infection across selection treatments for male and female mosquitoes, separately. We used a linear mixed model with the two forms of selection (reproduction and exposure to parasitism) as explanatory variables in interaction and replicate as a random factor. For fecundity, we assessed each window separately and quantified the number of eggs laid at 7, 21 and 35 days of adult age. For the best model fit, the number of eggs was transformed using a quadratic function. Using linear models similar to those for longevity, we quantified the cost of selection on reproductive life output.

| Tested effect | | | |
|---|---|---|---|
| **Model 1a**: Female longevity after selection | ***χ2*** | ***df*** | ***p*** |
| Selection reproduction | 20.62 | 1 | **< 0.001** |
| Selection parasitism | 8.07 | 1 | **0.005** |
| Selection reproduction : Selection parasitism | 2.09 | 1 | 0.148 |
| **Multiple comparisons for Model 1a** | **estimate** | ***t*** | ***p*** |
| Exposed x Early *vs* Naive x Early | 84.8 | 0.98 | 0.762 |
| Exposed x Early *vs* Exposed x Late | -362.7 | -4.25 | **0.013** |
| Exposed x Early *vs* Naive x Late | -101.8 | -1.19 | 0.652 |
| Naive x Early *vs* Exposed x Late | -447.6 | -5.17 | **0.004** |
| Naive x Early *vs* Naive x Late | -186.6 | -2.15 | 0.216 |
| Exposed x Late *vs* Naive x Early | 261.0 | 3.03 | 0.064 |
| **Model 1b**: Male longevity after selection | ***χ2*** | ***df*** | ***p*** |
| Selection reproduction | 1.29 | 1 | 0.260 |
| Selection parasitism | 4.23 | 1 | **0.040** |
| Selection reproduction: Selection parasitism | 0.11 | 1 | 0.738 |
| **Model 1cd:** Day 7 fecundity | ***χ2*** | ***df*** | ***p*** |
| Selection parasitism | 13.10 | 1 | **< 0.001** |
| Selection reproduction | 0.34 | 1 | 0.560 |
| Infection treatment | 0.44 | 1 | 0.508 |
| Selection parasitism : Selection reproduction | 8.01 | 1 | **0.005** |
| Selection parasitism : Infection treatment | 3.70 | 1 | 0.052 |
| Selection reproduction : Infection treatment | 0.35 | 1 | 0.556 |
| **Multiple comparisons for Model 1cd:** selection | **estimate** | ***t*** | ***p*** |
| Exposed x Early *vs* Naive x Early | 6.60 | 3.14 | **0.010** |

| | | | |
|---|---|---|---|
| Exposed x Early *vs* Exposed x Late | -0.61 | -0.29 | 0.991 |
| Exposed x Early *vs* Naive x Late | -2.49 | -1.17 | 0.644 |
| Naive x Early *vs* Exposed x Late | -7.21 | -3.39 | **0.004** |
| Naive x Early *vs* Naive x Late | -9.09 | -4.18 | **< 0.001** |
| Exposed x Late *vs* Naive x Early | -1.89 | -0.88 | 0.816 |
| **Model 1cd:** Day 21 fecundity | ***χ2*** | ***df*** | ***p*** |
| Selection parasitism | 0.11 | 1 | 0.741 |
| Selection reproduction | 5.84 | 1 | **0.016** |
| Infection treatment | 27.95 | 1 | **< 0.001** |
| Selection parasitism : Selection reproduction | 0.01 | 1 | 0.960 |
| Selection parasitism : Infection treatment | 0.81 | 1 | 0.368 |
| Selection reproduction : Infection treatment | 0.53 | 1 | 0.465 |
| **Model 1cd:** Day 35 fecundity | ***χ2*** | ***df*** | ***p*** |
| Selection parasitism | 18.40 | 1 | **< 0.001** |
| Selection reproduction | 5.43 | 1 | **0.020** |
| Infection treatment | 0.03 | 1 | 0.871 |
| Selection parasitism : Selection reproduction | 3.58 | 1 | 0.059 |
| Selection parasitism : Infection treatment | 2.27 | 1 | 1.132 |
| Selection reproduction : Infection treatment | 1.75 | 1 | 0.185 |

**Supplementary Table 2. MDA concentration across selection treatments.** We quantified oxidative damage by measuring malondialdehyde (MDA) concentration in adult female mosquitoes originating from lines selected for early or late reproduction, with or without exposure to parasitism. A linear model with reproduction- and parasite-selection as interacting fixed effects was used to test for differences in MDA levels. Shown are the Type II Wald $\chi^2$ statistics, degrees of freedom, and *p*-values for each fixed effect in Model 2a.

| **Tested effect** | | | |
|---|---|---|---|
| **Model 2a** | ***χ2*** | ***df*** | ***p*** |
| Selection reproduction | 1.33 | 1 | 0.248 |
| Selection parasitism | 5.16 | 1 | **0.023** |
| Selection reproduction : Selection parasitism | 4.63 | 1 | **0.031** |

**Supplementary Table 3. Summary of the principal component analysis.** PCA variance in the presented Figure 2b (underlined values). Eigenvalues, proportion of variance, and cumulative variance explained by the first principal components describing variation in oxidative stress markers across selection lines. These axes were used to summarise the multivariate redox gradient among lines selected for early or late reproduction, with or without exposure to parasitism.

| | **Eigenvalue** | **Proportion of variance** | **Cumulative variance** |
|---|---|---|---|
| **PC1** | 1.955 | 0.391 | 0.391 |
| **PC2** | 1.659 | 0.332 | 0.723 |
| **PC3** | 1.025 | 0.204 | 0.927 |
| **PC4** | 0.236 | 0.047 | 0.975 |
| **PC5** | 0.125 | 0.025 | 1.000 |

**Supplementary Table 4. Loadings of oxidative markers on PCA.** Loadings of mean GSH, GSSG, redox ratio, MDA, and SOD concentrations on the first two principal components from the PCA of oxidative markers. Positive and negative values indicate the direction and strength of each trait's contribution to the multivariate redox gradient separating selection treatments.

| Trait | Loading PC1 | Loading PC2 |
|---|---|---|
| **GSH/GSSG ratio** | 0.634 | -0.281 |
| **GSSG** | 0.535 | -0.418 |
| **GSH** | -0.455 | -0.499 |
| **SOD** | -0.308 | -0.652 |
| **MDA** | 0.096 | -0.268 |

**Supplementary Table 5. Relationship between oxidative PCA axis and longevity**. Linear (LM) and generalized additive (GAM) models relating median adult longevity (uninfected assay) to PC1 scores from the PCA of oxidative markers (**Figure 2b,c; Supplementary Tables 3 and 4)**. For the LM, slope estimate, standard error, t-value, p-value, $R^2$ and adjusted $R^2$ are shown. For the GAM, effective and reference degrees of freedom (edf, ref. df), F-value, p-value, $R^2$ and explained deviance are shown. Negative slopes indicate that lines with more oxidized redox profiles (higher PC1 scores) tend to have reduced median lifespan.

| | **estimate** | **std. error** | ***t*-value** | ***p*-value** | **$R^2$** | **Adjusted $R^2$** |
|---|---|---|---|---|---|---|
| **PC1 (LM)** | -0.876 | 0.404 | -2.168 | 0.055 | 0.320 | 0.252 |
| | **edf** | **ref. df** | ***F*-value** | ***p*-value** | **$R^2$** | **Explained dev.** |
| **PC1 (GAM)** | 2.18 | 2.68 | 5.00 | **0.025** | 0.551 | 0.64 |

### Figures

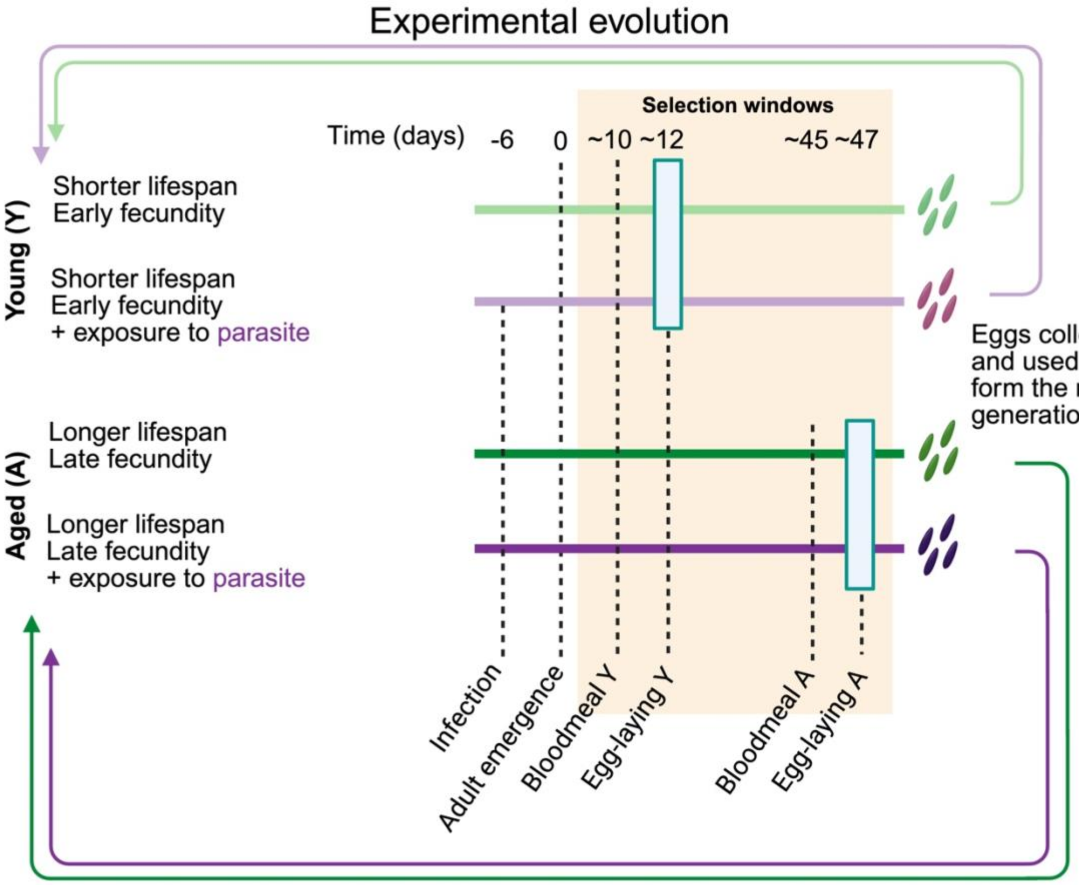


**Supplementary Figure 1. Design of the experimental evolution setup.** Illustration of the experimental evolution assay. From a single starting stock population of *Aedes aegypti*, four selection treatments were created: (i) Exposed to parasite x Early reproduction; (ii) Exposed to parasite x Late reproduction; (iii) Naive x Early reproduction; and (iv) Naive x Late reproduction. Each selection treatment is composed of four replicate lines run in parallel. Lines selected for early reproduction had the offspring from the first blood meal selected to start the next generation. In contrast, the late-reproduction lines were blood-fed weekly, and the selected clutch of eggs was the result of the blood meal around 45 days after adulthood (when approximately two-thirds of the population had died). Lines exposed to the parasite *V. culicis* were subjected to an infective dose of 20,000 spores per larva, 2 days after eggs had hatched. This was repeated for at least ten generations before measuring all the traits.

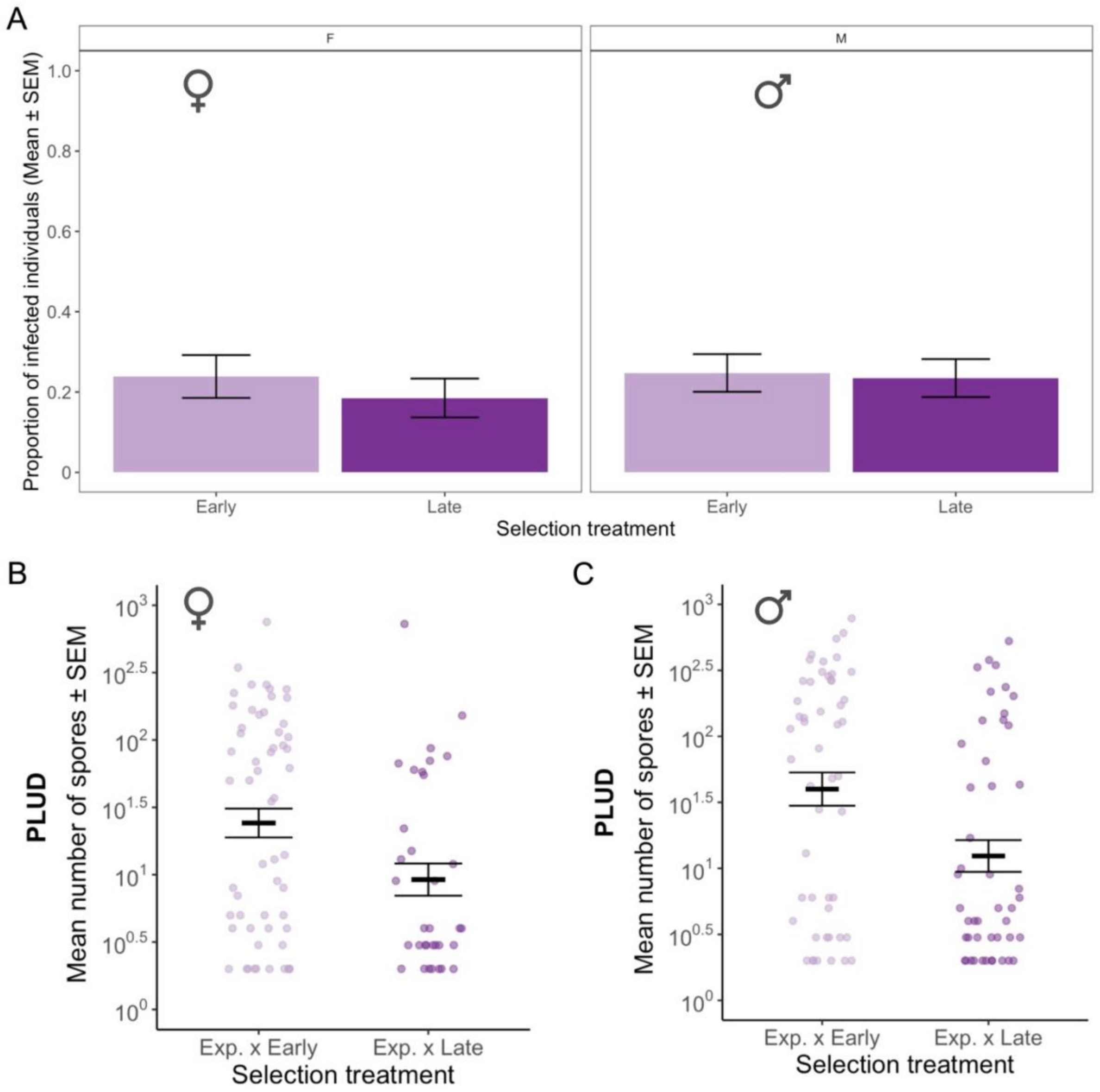


**Supplementary Figure 2. Post selection measurements: parasite load at death.** As individuals died during the longevity assay, they were collected to quantify **(A)** the proportion of infected females and males. From those who did not clear infection, the mean number of spores was counted and compared for **(B)** females, and **(C)** males. No statistical difference was found between treatments for any of the comparisons.

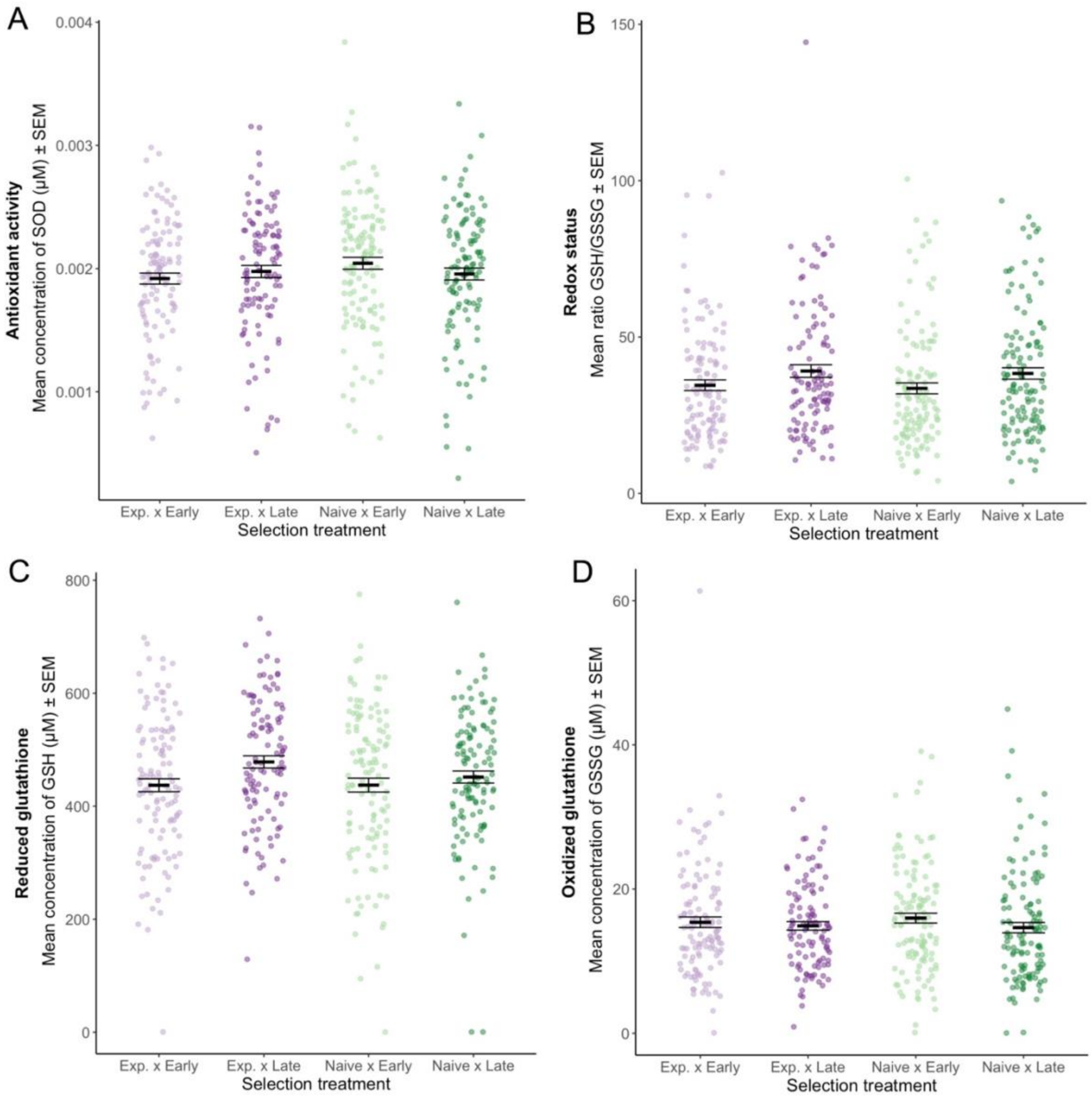


**Supplementary Figure 3. Post selection measurements: oxidative markers.** Mean concentrations, and respective errors, of **(A)** superoxide dismutase (SOD, marker of antioxidant activity), **(B)** the GSH: GSSG ratio (proxy for redox status), **(C)** reduced glutathione (GSH), and **(D)** oxidized glutathione (GSSG) measured in uninfected females from each selection treatment at day 5 and 15 of age. There was no difference across selection treatments in either time point, so they were combined. None of the markers showed any difference across selection treatments.
